# Monte Carlo Simulation Analysis of Variant Burden Prompts Potential Oligogenic Interactions

**DOI:** 10.64898/2025.12.27.696666

**Authors:** Villem Pata, Markus Marandi, Jaan Märten Huik, Katrin Õunap, Sander Pajusalu

**Affiliations:** Department of Genetics and Personalised Medicine, Institute of Clinical Medicine, University of Tartu, Tartu, Estonia; Genetics and Personalized Medicine Clinic, Tartu University Hospital, Tartu, Estonia; Anesthesiology and Intensive Care Clinic, Tartu University Hospital, Tartu, Estonia; Institute of Computer Science, University of Tartu, Tartu, Estonia; School of Engineering Sciences in Chemistry, Biotechnology and Health, KTH Royal Institute of Technology, Stockholm, Sweden; Department of Radiology, Institute of Clinical Medicine, University of Tartu, Tartu, Estonia

## Abstract

**Background:** A large fraction of patients with suspected rare genetic disorders remain undiagnosed, suggesting that complex inheritance patterns, which are poorly detected by standard monogenic tests, may be a key factor.

**Methods:** We developed a computational framework that uses Monte Carlo simulations on summary-level allele counts to test for pathway-level burden enrichment. This approach was applied to a cohort of 639 neuromuscular disease (NMD) patients and 9,059 controls from the Tartu University Hospital (2015-2023) as proof of concept and compared with standard burden testing.

**Results:** While standard per-gene Z-tests failed to yield genome-wide significant results, our simulation framework identified a nominal enrichment of rare, moderate-impact variants within a predefined set of NMD-associated genes (Bonferroni-corrected empirical p = 0.02). Exploratory analysis of the most significantly burdened gene sets revealed frequent co-occurrence of *CAPN3* with other critical NMD genes, including *CLCN1, GFPT1*, and *MYOT*.

**Conclusion:** Our findings suggest a potential oligogenic model in NMD, where variants in a hub gene like *CAPN3* may interact with other rare variants to contribute to disease. This Monte Carlo simulation framework needs further validation but demonstrates that this framework may be a useful tool for hypothesis generation in complex genetic disorders.

**Funding:** Estonian Research Council grants PSG774 and PRG2040.

**Availability:** publicly available under CC BY-NC-SA 4.0 licence at https://github.com/OligoGeneticDiseases/gen-toolbox

## Introduction

Despite ever-increasing Next Generation Sequencing (NGS) data, growing databases, advanced variant detection and classification methods and rapidly increasing knowledge from research studies, the diagnostic yield in patients with suspected genetic disorder remains below 50% (1–4). One of the proposed explanations to relatively low diagnostic sensitivity are more complex non-Mendelian modes of inheritance, such as digenic or oligogenic interactions (1,2). Databases and tools like DIDA, Orval, VarCoPP2.0, Hop and DiGePred have provided new evidence and clinical utility for digenic interactions with the use of machine learning(1–4) — often relying on known digenic interactions. Elucidating complex interactions of rare variants in different gene pathways remains challenging.

Variant burden analysis is one group of statistical methodologies that aggregates a group of rare variants in a given locus such as a gene and regresses against a phenotype to show the cumulative effect of the rare variants in the region (e.g. cohort allelic sums test, CAST)(4). Early variant burden tests grouped variants with different functional effects, while later tests added variant “weights” based on functional effect scores or allele frequency to increase power, Boutry et al presenting a classification for available tools (5). Using allele frequency as a proxy for causality stems from the idea that alleles with a selective disadvantage have low frequency and remain infrequent due to purifying effect (1,2,5). The usefulness of burden analysis in correlating rare variant burden to phenotype in both simulated and empirical data has been shown by earlier authors (1,2,5). Price et al. proposed using resequenced allele data and pooling alleles of complete gene pathways to reduce Type I error and hence missed cases (6).

Building on this pathway-level logic, several computational frameworks have been developed to assess aggregate enrichment. Pascal (Pathway Scoring Algorithm) (7) utilizes an analytical approach to convert gene-level p-values into a pathway score. Alternatively, aSPUpath (Adaptive Sum of Powered Score) (8) employs an adaptive weighting scheme for rare variants, using Monte Carlo simulations to calculate empirical significance. Similarly, MEGA-V (Gene set enrichment analysis for rare variants) (9) uses phenotype permutations to evaluate the significance of variant burdens within specific gene sets.

Our proposed statistical methodology differs from these established tools by operating on summary-level allele count matrices rather than individual-level genotypes, the summary matrices can be generated with the accompanying pipeline. Furthermore, we utilize a non-parametric Monte Carlo (10,11) simulation framework to generate and compare empirical distributions of burden ratios generated from cases and controls. Further ontological analysis is done to correlate these genes to the proposed phenotype.

Our primary hypothesis is that an oligogenic architecture can be detected as a statistical shift in the distributions of variants when stratified by rarity and functional impact. Under this model, ultra-rare, high-impact variants, such as protein-truncating alleles, would typically correlate with a classical monogenic burden. In contrast, we propose that more complex phenotypes are driven by an accumulation of non-loss-of-function (non-LoF) variants, which are individually less deleterious but collectively exceed a physiological threshold for disease when present in specific pathways. To demonstrate this methodology for oligogenic burden analysis, we use an example clinical cohort of patients with neuromuscular disease (NMD), both with and without molecular diagnosis. While we demonstrate proof of concept on targeted gene panels, the same framework extends to whole exome and genome sequencing data.

## 2. Materials and Methods

### 2.1 Study Population and Cohort Definition

#### 2.1.1 Clinical Cohort and Phenotyping

The study utilized a clinical cohort from Tartu University Hospital (2015–2022) consisting of 9,746 individuals. After quality control 9,698 samples were retained. Ethics approval was granted by the University of Tartu Research Ethics Committee for reanalysis of variant files. In accordance with the research permit 374/M-6, variant data contained in VCFs can only be correlated with only very coarse anonymous information available (referral, annotation metadata, date). Phenotype data is based on the original clinical referral presented to the testing lab and were additionally manually curated by a clinician. The phenotype-positive group comprised 639 patients referred for neuromuscular disease (NMD), manually curated by clinical specialists. Where individuals presented with several phenotypes, NMD was selected as the primary phenotype.

#### 2.1.2 Selection of the Null Cohort

The remaining 9,059 individuals referred for non-NMD indications such as intellectual disability, epilepsy, metabolic disorders, congenital heart disease, connective tissue disorders, healthy individuals sequenced after genetic counselling for family planning, etc served as the control pool. To address potential recruitment bias, we generated a “null cohort” by randomly selecting 639 samples from this pool to serve as a size- and sex-matched comparison group for global distribution testing (see Supplementary Data, Figure 1 and Table 2).

**Figure 1.**
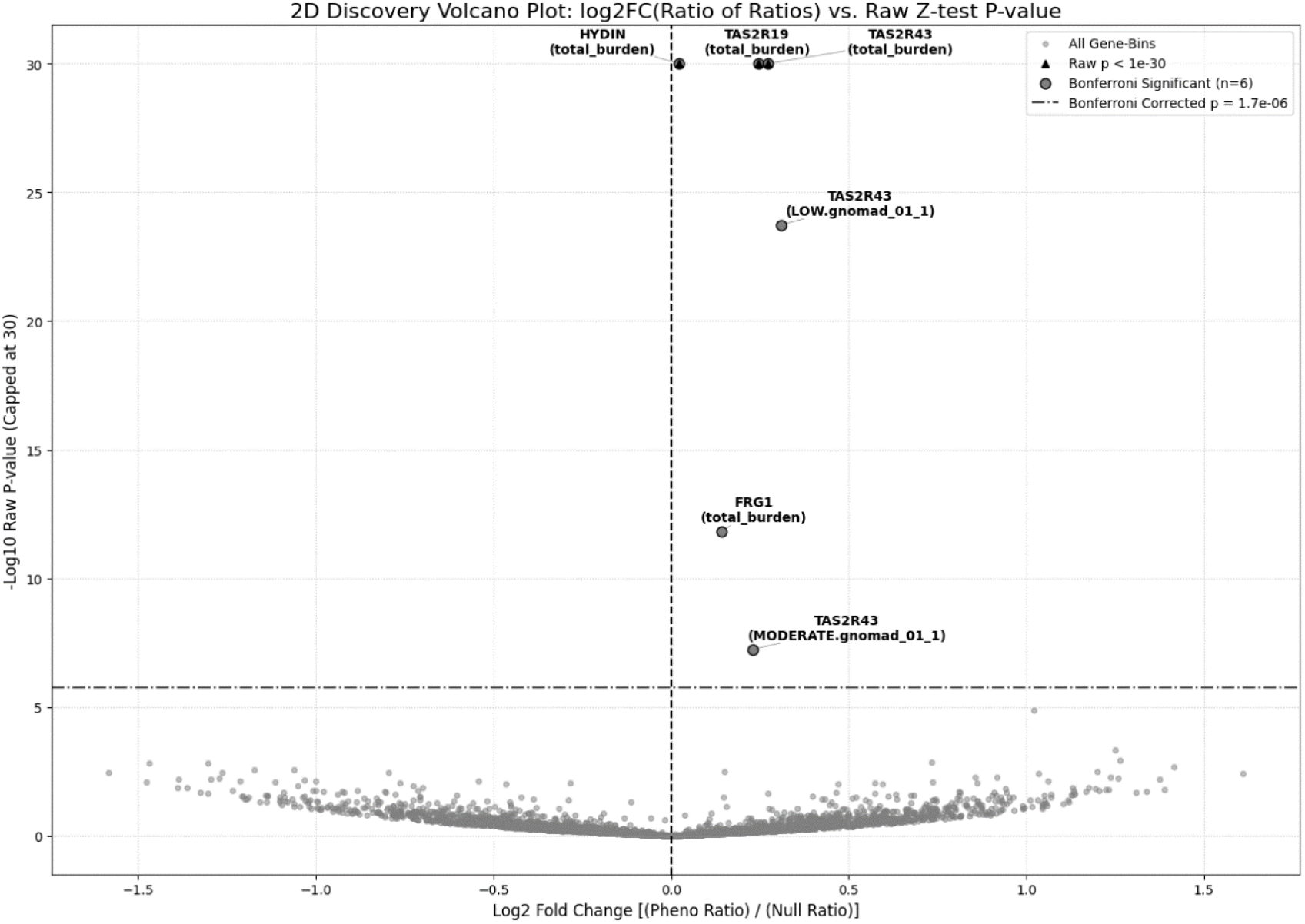
Volcano plot identifying gene-bins with significant differential burden between phenotype and null cohorts. Each point represents one of 29,801 gene-bin tests. The y-axis displays statistical significance as the negative log10 of the raw p-value, derived from a Z-test comparing the log-odds ratios of allele burden between the phenotype and null cohorts where able to calculate (i.e., allele count > 1 in both cohorts). The x-axis represents the effect size, calculated as the log2 fold change of the case-control burden ratios (Phenotype Ratio / Null Ratio). The horizontal dashed line indicates the Bonferroni-corrected significance threshold (p < 1.7×^-6^). Labelled are six highly significant gene-bins (e.g., HYDIN, TAS2R19, FRG) with high statistical confidence but a log2 fold change near zero, suggesting that their magnitude relative to the null cohort is small, the small p-values arising from the relatively larger allele counts compared to most datapoints.

## 3. Genetic Data Processing and Quality Control

Samples were sequenced using targeted panels (Illumina TruSight One or TruSight One Expanded), covering an intersection of 4,481 genes. Reads were aligned to the GRCh37 reference genome, with the variant calling that had already passed quality analysis in the original clinical diagnostic pipeline. We also successfully created summary burden statistics with our Illumina NovaSeq Xplus-sequenced DRAGEN-annotated whole exome dataset, but these samples were excluded from analysis due to the large imbalance in panel sizes.

Relationship inference was performed using KING-estimated kinship coefficients(12). We applied a permissive threshold of >0.354 to exclude only monozygotic twins and duplicates. This threshold was chosen to retain first-degree relatives, which act as biological replicates in rare-variant studies for low-prevalence disorders. Our proof-of-concept dataset relies on targeted gene panel data, where the sparsity of common markers (restricted to panel targets) increases the variance of kinship estimates, posing a risk of excluding unrelated individuals (false positives) at lower thresholds. As such, 48 samples were pruned during quality control due to likely repeat sequencing of individuals with new unique IDs.

The ancestry of the enrolled patients was almost homogenously reported to be Estonian (EUR) ancestry. Sample sex was inferred from the binomial clustering of the F-statistic calculated from variants on the X-chromosome, excluding the pseudoautosomal region (vcftools (13)). A chi-square analysis confirmed that there was no significant sex distribution bias between the neuromuscular disease (NMD) and null cohorts (χ^2^=1.35 (p=0.25)).

## 4. Variant Annotation and Categorization

The toolkit aggregates variants into groups by Gene Label, Functional Impact and Minor Allele Frequency (MAF) with all variants having equal weight within their respective group. Variants were reannotated using Ensembl Variant Effect Predictor (VEP v111) (14) for cohesion. Gene labels corresponded to HUGO Gene Nomenclature Committee (HGNC) labels covered by the targeted sequencing panels and were also VEP-annotated. VEP additionally categorized variants into 4 Functional Impact tiers based on transcript Sequence Ontology terms, analysis was narrowed down to High (protein-truncating variants e.g., stop-gained, frameshift), Moderate (in-frame indels and missense variants), Low impact tiers (Synonymous and non-canonical splice region variants) (comprehensive categorization in Table 1, Supplementary Data). This type of categorization was selected for the low computational cost, however the end-user may implement other VEP-annotated prediction plugins i.e. AlphaMissense (15) or MPC scores (16) for pathogenicity tiering.

Minor Allele Frequencies (MAF) were derived from gnomAD (17). Variants were binned into three frequency ranges: <0.01%, 0.01-0.1%, 0.1-1% (3). Thus, common variants were excluded from burden aggregation.

The aggregated minor allele counts (MAC) for both hetero- and homozygous variants in nine bins (3x3) + the total allele counts for every gene were used for downstream analysis between phenotype positive and negative cohorts. This grouping of variants based on allele frequency and transcript consequence ensures any variant may be thus classified and is not computationally intensive to annotate even on biobank-level datasets. It is expected that such categorization of variants may produce low MACs in some bins, i.e., it is unlikely to find more common but also very deleterious protein truncating variants.

## 4. Statistical Framework for Burden Analysis

### 4.1 Summary-Level Matrix Generation

Aggregation of variants by gene, impact, MAF into the 10 bins was done using our Hail-based (Hail v0.2.133 https://github.com/hail-is/hail)) toolkit, which is scalable from a local machine to cloud hosting services. The aggregated summary matrices are then used for statistical analysis. This approach of using summary-level data for statistical analysis from locally generated datasets enables data sharing and collating of data from restricted biobanks but has limitations based on data granularity.

### 4.2 Single-Gene, Global Distribution and Monte Carlo simulations

We applied a one-sided Z-test for difference in proportions to evaluate single-locus burden enrichment within the NMD cohort. To account for the sparse, zero-inflated allele counts typical of rare variant subgroups, we implemented the Haldane-Anscombe correction (18,19), adding a +0.5 continuity adjustment to all observed counts. This adjustment ensures defined variance estimates and minimizes bias in standard error computation when allele counts are low, enabling stable inference where traditional asymptotic approximations fail. To focus the analysis on informative loci, gene-bin combinations were only included if the sum of allele counts in both the case and control groups was ≥ 1 for both the phenotype and null cohorts. To maintain a stringent family-wise error rate, Bonferroni correction was applied based on the total number of gene-bin tests performed (n=29,801, α=1.7 × 10^-6^).

### 4.3 Pathway-Level Enrichment

To detect additive oligogenic signals, we implemented a non-parametric Monte Carlo simulation framework. A single simulation is defined as the random selection of 10 genes to form a “synthetic pathway.” For each pathway, we calculate the burden ratio (sum of minor alleles in cases divided by the sum in controls). Due to the sparsity of rare variant data, we only included simulations where the aggregate MAC for the 10-gene set was ≥ 1. This permissive filter ensures that “singleton” variants are retained, avoiding the loss of signal associated with more stringent frequency filters. To prevent division by zero, a small constant (ε = 10×^-9^) was added to all aggregate burdens.

To maximize sensitivity and assess biological specificity, this process was repeated for 10^6^ Monte Carlo simulations per categorical bin across two gene pools:

1. Hypothesis-free Discovery: Gene sets were drawn from the entire available pool of ∼4,500 genes. This serves as a global baseline, representing the general background of rare variation across the targeted panels.
2. A predefined NMD phenotype (Genomics England PanelApp Level 3: Neuromuscular Disorders) (20), narrowing the potential interactions down to known biological pathways.

We utilized a dual-stage statistical evaluation. First, penalized Firth’s Logistic Regression (21,22) with (PyPI firthlogist v0.5) (23) was used for a global comparison of the cohort distributions to mitigate bias in sparse datasets, addressing the question: “is there shift in global enrichment?”

Second, empirical p-values were calculated based on the frequency with which the null cohort’s ratios exceeded the 99^th^ percentile (tail) of the phenotype cohort’s distribution, addressing: “is there a tail-end enrichment in burden?”

The resulting empirical tail-end p-values were Bonferroni-adjusted for 20 independent tests (10 bins ×2 sampling pools). By comparing these two pools, we can statistically differentiate between general variant accumulation and phenotype-specific enrichment. This serves as an internal control: significant tail-end enrichment in NMD-targeted sets over the global background strengthens the evidence for biological interactions over cohort-specific bias.

### 4.4 Identification of Hub Genes

To dissect the specific drivers of the observed burden in the category shown to have tail-end statistical enrichment between the NMD and Null cohorts, we performed a pairwise co-occurrence analysis. This analysis proceeded in three steps: (1) we identified the top 25 contributing genes based on their frequency of appearance within the top 1st percentile (99^th^ percentile tail) of all simulated gene sets; (2) we calculated pairwise co-occurrence counts for these genes; and (3) we ranked these pairs to identify ‘Hub’ genes that consistently anchored high-burden sets. Frequently appearing genes and pairs would be used for hypothesis-generation and, later, more granular, sample-specific analysis.

## 5. Code and Data Availability

The Hail-based toolkit for oligogenic analysis and statistical analysis pipeline is available under the Creative Commons Attribution-NonCommercial-ShareAlike 4.0 International Public License. The toolkit can be publicly accessed at https://github.com/OligoGeneticDiseases/gen-toolbox. Original individual VCF data is not publicly available due to data restrictions on sensitive clinical information imposed by numerous national, EU and hospital regulations. Summary anonymous variant data collated from individual data using the Hail-based tool can be accessed on Github. The statistical analysis is provided in a Jupyter notebook. Data analysis was done on an instance of Tartu University High Performance Computing Center cluster (24). Funding was provided by Estonian Research Council grants PSG774 and PRG2040.

## 6. Results

### 6.1 Single-Gene Association Tests Yield Limited Findings

The one-sided Z-test yielded for standard single gene-based burden comparing the cumulative frequency of rare variants (MAF < 1%) between the NMD (N=639) and control cohorts (N=9,059) identified four unique loci with significant difference in odds-ratios after Bonferroni correction: (p < 1.7×10^-6^ for 29801 tests) for *HYDIN, TAS2R19, TAS2R43* and *FRG1* (Figure 1).

The limited number of findings from this method underscores the difficulty in detecting signals for disorders with complex inheritance patterns using just single-gene approaches and motivates the search for oligogenic patterns.

### 6.2. Hypothesis-Driven Simulation Uncovers Significant Burden in NMD-Specific Gene Pathways

For Monte Carlo simulation analysis we generated one million random 10-gene sets from all available genes in the intersect of the summary MAC table for each of the 10 available categorical bins. Firth’s logistic regression per bin showed very small but statistically significant differences (OR=0.9 … 1.1, p<1.5×^-10^) in distributions between the two cohorts (see Supplementary Data, Table 3), likely owing to the very large number of datapoints available for regression.

To test the specific hypothesis that a cumulative burden of rare variants exists within known NMD pathways, the one-sided empirical p-value was calculated for all genes vs NMD genes. A comparison of the aggregate burden scores from these NMD-specific sets revealed a significant rightward shift in the distribution for the NMD cohort versus the Null cohort, confirming an enrichment of rare variants within these pathways (Figures 2 & 3). The signal was most pronounced for variants classified as having a moderate functional impact and a MAF < 0.01% (‘MODERATE.gnomad_001’ bin), which showed a significant burden increase (empirical p = 0.02, Bonferroni corrected, 20 tests) and low impact, MAF 0.01% to 0.1% (‘LOW.gnomad_001_01’) bin. We focused on the moderate category per our original hypothesis for oligogenic effect appearing as rare but non-LoF variants.

To identify which specific genes within this curated NMD set were the primary drivers of the signal in the ‘MODERATE.gnomad_001 bin’, we analysed the composition of the gene sets from the top percentile of the burden distribution. This analysis identified a ranked list of “top contributing” genes: *MYOT, MYF6, ETFA, TK2, FKRP, SLC25A4, GFPT1, ALG2, CAPN3, CFL2, TCAP, FKBP14, TPM2, DPAGT1, CLCN1, TNNI2, CYP2C8, TSEN54, DES, ACTA1, FHL1, SLC5A7, LAMP2, HRAS* and *FKTN*.

Although we hypothesized that an increased burden of rare high impact variants would be present in the NMD cohort, our simulation framework did not detect a statistically significant enrichment with a set size of 10 genes.

**Monte Carlo simulation analysis Figures 2 and 3.**

**Figure 2.**
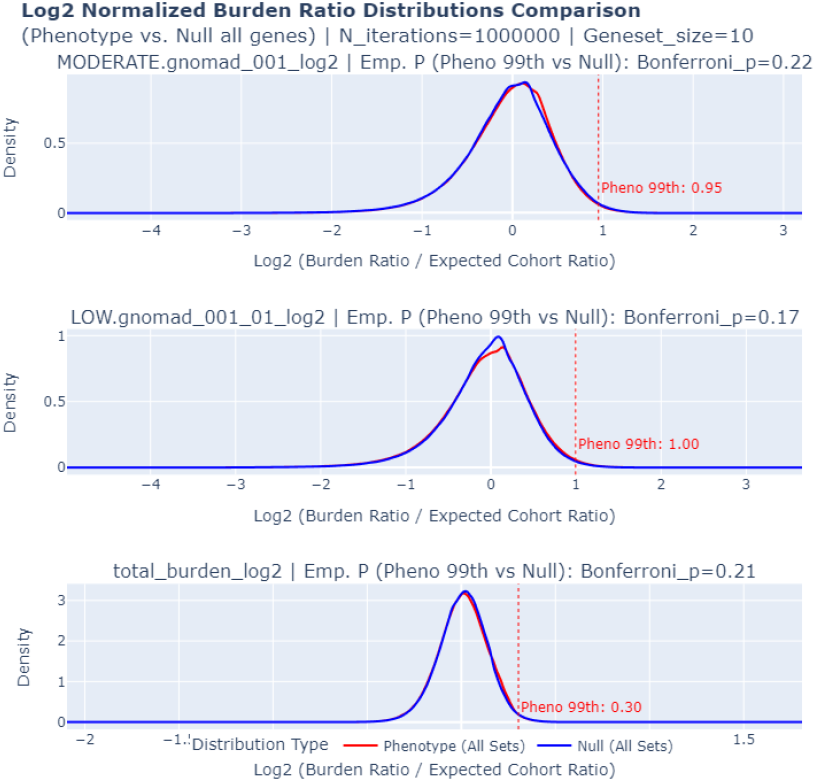
Gene sets from all genes. Only 3 bins out of 10 shown. Monte Carlo simulations of 10 genes over 10 bins, 1e6 simulations per bin. Empirical p-values for the 99th percentile were derived from phenotype (red) to null (blue) cohort comparison. No high tail-end burden apparent as p>0.05 after Bonferroni correction in all bins. “HIGH” bins are non-normal due to the smaller number of burdened genes. Truncated (see Supplementary Data).

**Figure 3.**
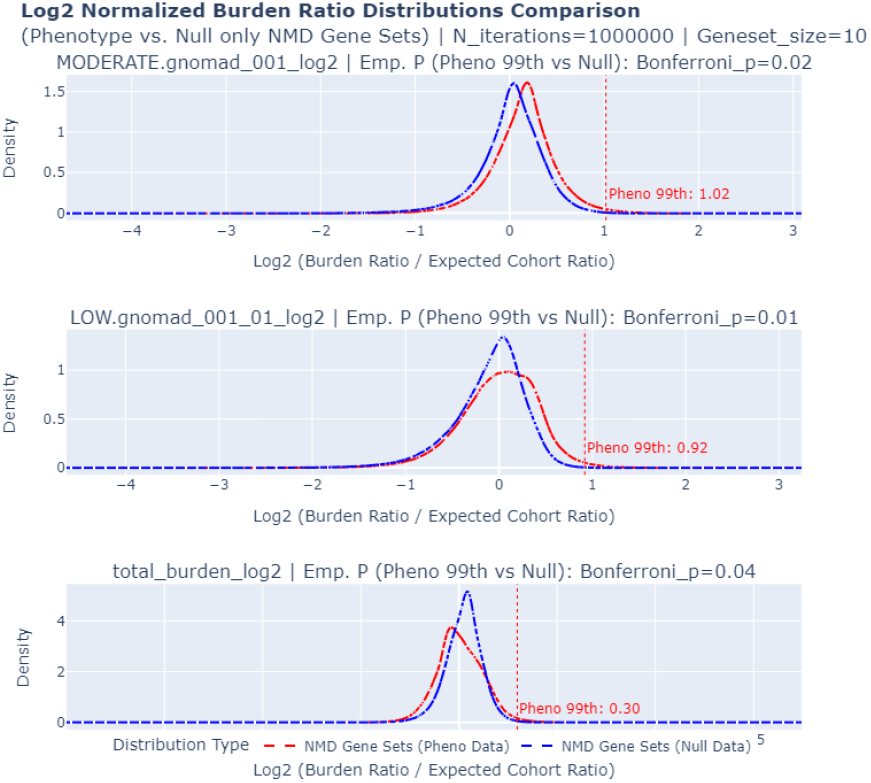
Gene sets from Neuromuscular Disorders gene panel. Monte Carlo simulations of 10 genes in 10 bins, 1e6 simulations per bin. Empirical p-values for the 99th percentile were derived from phenotype (red) to null (blue) cohort comparison. High tail-end burden increase was detected in moderate < 0.001%, low 0.001…0.01% and total burden bins. Truncated (see Supplementary Data).

### 6.3 Co-occurrence Analysis Reveals *CAPN3* as a Central Hub in a Potential Oligogenic Network

The pairwise co-occurrence analysis identified the Calpain-3 gene (*CAPN3*) as a central interaction hub, specifically within the ‘Moderate Impact’ (MAF < 0.01%) bin based on the most pronounced signal in the high-tail end burden in when comparing the

The most frequent interacting partners for *CAPN3* included *CLCN1* (associated with myotonia congenita), *GFPT1* (congenital myasthenic syndrome), *FKRP* (limb-girdle muscular dystrophy), and *MYOT* (myofibrillar myopathy). KEGG (25) pathway analysis further supported a biological basis for our findings as eight of the top 25 genes mapped to “Cytoskeleton in muscle cells” (hsa04820), seven to “Metabolic Pathways” (hsa01100), three to “Motor proteins” (hsa04814) illustrated in Figure 4.

**Figure 4.**
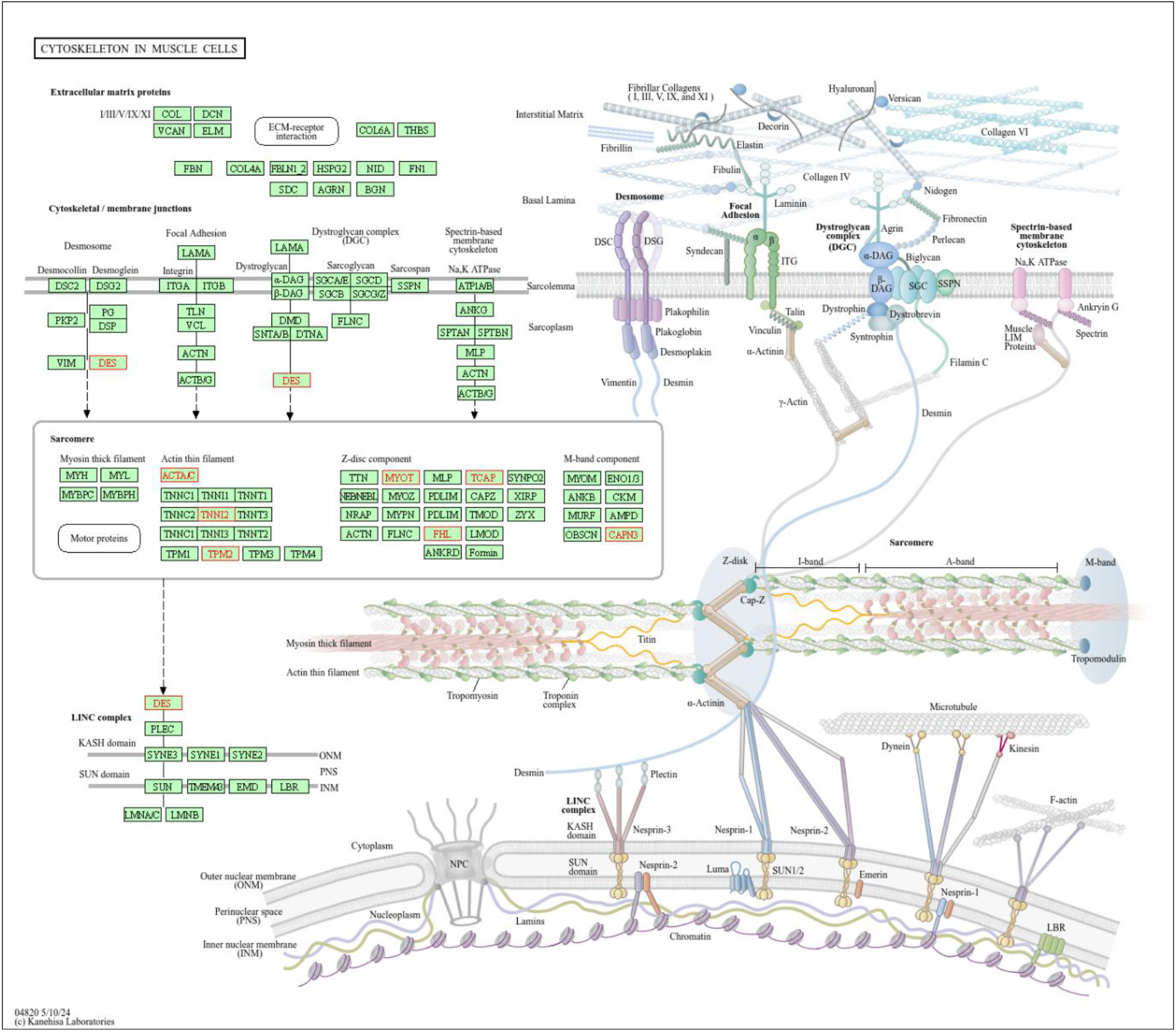
KEGG pathway for Cytoskeleton in muscle cells. Significant genes in red. “Cytoskeleton in muscle cells” (hsa04820), 7 to “Metabolic Pathways” (hsa01100), 3 to “Motor proteins” (hsa04814). Generated on the KEGG platform. Publishing dependent on Copyright Permission, requested from Kanehisa Laboratories.

## 7. Discussion

### 7.1 Hypothesis generation as primary benefit

In this study, we present a computational framework for detecting oligogenic burden signals in large-scale sequencing data using summary-level variant counts. The diagnostic gap in rare genetic disorders—frequently exceeding 50%—has motivated the search for complex inheritance models.

We implemented two complementary approaches to capture distinct genetic signals. While the Z-test with Haldane–Anscombe correction was optimized for single-locus, sparse, zero-inflated allele counts, the Monte Carlo framework was designed to assess pathway-level enrichment without relying on parametric assumptions. The primary innovation in our simulation framework is the use of a dual-cohort empirical distribution comparison. If gene sets were sampled only from the “all genes” pool within the phenotype cohort, the resulting empirical p-value would lack a biological baseline to account for potential systematic biases, such as sequencing batch effects or population-specific rare variant inflation. To address this, we generated parallel burden ratio distributions for both the NMD cohort and a size-matched Null cohort. The “burden ratio” for any given gene set was defined as the ratio of observed allele counts to the expected cohort ratio. By testing the null hypothesis that the NMD cohort’s distribution is not significantly shifted toward higher values compared to the null cohort’s distribution, we effectively controlled for the background “noise” of rare variation. This comparative approach enables the detection of a true biological signal—an enrichment of rare variants specifically within NMD-associated pathways—that would otherwise be indistinguishable from the heavy-tailed background distribution of rare variants across the genome. Limiting to a certain gene set reduces the ability to find completely new pathways but helps concentrate on the burden of combinations of loci that correlate to the phenotype.

We observed no significant concordance between the rankings obtained from the Z-test and the simulation framework. Notably, the known neuromuscular disease gene *CAPN3* fell below significance thresholds in the standard Z-test but was identified as a central “Hub” by the Monte Carlo co-occurrence analysis. Crucially, this finding emerged from the ‘Moderate Impact’ bin (MAF < 0.01%), which excluded high-impact protein-truncating variants. While this exclusion reduces the influence of obvious null alleles, we acknowledge that the *CAPN3* signal likely represents a composite of both oligogenic modulators and cryptic monogenic drivers. The primary utility of this framework is its ability to prioritize such hubs agnostically: whether the underlying mechanism is strictly oligogenic or a monogenic burden, the tool successfully flags a relevant NMD driver gene for downstream segregation analysis.

Our analysis prompted multiple distinct pathways, likely reflecting the phenotypic heterogeneity of the broad NMD cohort. While stricter phenotypic grouping might isolate single pathways, it comes at the cost of reduced sample size and statistical power. We also observed that strong monogenic signals may be diluted as simulation set sizes increase, representing a trade-off between detecting complex interactions and identifying high-effect Mendelian variants.

A defining feature of our approach is its privacy-preserving, customisable, scalable workflow. Built on the Hail toolkit, our pipeline scales from single machines to distributed computing clusters, enabling the ingestion of large datasets—such as Illumina DRAGEN output or biobank-level VCFs—to generate anonymous summary matrices. In addition, unlike statistical methods requiring individual-level genotypes throughout (e.g., SKAT), our statistical analysis operates exclusively on these summary matrices which can be additively collated. This decoupling enables the retrospective aggregation of sensitive clinical cohorts across institutions and sequencing platforms without sharing individual patient data. The authors have chosen to focus on the well-defined neuromuscular pathway; however, the toolkit enables researchers to explore other phenotypes and fuzzier hypothetical pathways.

### 7.2 Limitations

Our methodology is not immediately applicable to increase the diagnostic yield in the clinic. Several limitations must be considered before taking this proof-of-concept analysis at face value.

The individuals in our control pool were referred for clinical genetic testing for non-neuromuscular indications. While size-matched for the null cohort, these individuals are not all healthy controls in the traditional sense, and it is possible that overlapping sub-clinical phenotypes or unrelated genetic burdens could influence the background distribution (e.g. connective tissue disorders and neuromuscular disease may be difficult to clinically differentiate).

In this study, we use “interaction” to describe the cumulative or additive burden of rare variants within a biological pathway or gene set. Our current summary-level approach does not explicitly model second-order epistatic (G×G) interactions or suppressive gene interactions.

We observed that known neuromuscular disease genes, such as *CAPN3*, acted as central hubs in our co-occurrence analysis. While we restricted the primary analysis to ‘Moderate Impact’ variants to minimize obvious null alleles, the signal likely represents a composite of both potential oligogenic modulators and some monogenic drivers.

Our NMD cohort represents an undifferentiated spectrum of neuromuscular disorders. While this increased our sample size, the high degree of phenotypic heterogeneity may have diluted signals from specific pathways that might be more apparent in more strictly defined clinical subgroups. Finally, we have not provided a benchmark comparison with existing statistical tools.

## 8. Conclusion

Overall, our results suggest that the Monte Carlo simulation-based statistical methods provide an alternative to conventional rare variant tests for detecting oligogenic burden. Specifically, discovery sets of potential oligogenic pathways can be created in the high tail-end of variant burden and we showed that these pathways correlate to our example neuromuscular disease cohort with KEGG pathway analysis. The results are limited in detecting specific variant combinations due to the focus being on summary level data.

Further correlation needs to be done on a per-individual and per-variant testing using already available oligogenic datasets for validation, i.e. Orval and DIDA; integrating AlphaMissense predictions for more accurate pathogenicity tiering and doing phenotype-permutation validation with MEGA-V.

