## Supplementary Data for "Monte Carlo Simulation Analysis of Variant Burden Prompts Potential Oligogenic Interactions"

| SO term | Description | Display term | IMPACT |
|---|---|---|---|
| transcript_ablation | A feature ablation whereby the deleted region includes a transcript feature | Transcript ablation | HIGH |
| splice_acceptor_variant | A splice variant that changes the 2 base region at the 3' end of an intron | Splice acceptor variant | HIGH |
| splice_donor_variant | A splice variant that changes the 2 base region at the 5' end of an intron | Splice donor variant | HIGH |
| stop_gained | A sequence variant whereby at least one base of a codon is changed, resulting in a premature stop codon, leading to a shortened transcript | Stop gained | HIGH |
| frameshift_variant | A sequence variant which causes a disruption of the translational reading frame, because the number of nucleotides inserted or deleted is not a multiple of three | Frameshift variant | HIGH |
| stop_lost | A sequence variant where at least one base of the terminator codon (stop) is changed, resulting in an elongated transcript | Stop lost | HIGH |
| start_lost | A codon variant that changes at least one base of the canonical start codon | Start lost | HIGH |
| transcript_amplification | A feature amplification of a region containing a transcript | Transcript amplification | HIGH |
| feature_elongation | A sequence variant that causes the extension of a genomic feature, with regard to the reference sequence | Feature elongation | HIGH |
| feature_truncation | A sequence variant that causes the reduction of a genomic feature, with regard to the reference sequence | Feature truncation | HIGH |
| inframe_insertion | An inframe non synonymous variant that inserts bases into in the coding sequence | Inframe insertion | MODERATE |
| inframe_deletion | An inframe non synonymous variant that deletes bases from the coding sequence | Inframe deletion | MODERATE |
| missense_variant | A sequence variant, that changes one or more bases, resulting in a different amino acid sequence but where the length is preserved | Missense variant | MODERATE |
| protein_altering_variant | A sequence_variant which is predicted to change the protein encoded in the coding sequence | Protein altering variant | MODERATE |
| splice_donor_5th_base_variant | A sequence variant that causes a change at the 5th base pair after the start of the intron in the orientation of the transcript | Splice donor 5th base variant | LOW |
| splice_region_variant | A sequence variant in which a change has occurred within the region of the splice site, either within 1-3 bases of the exon or 3-8 bases of the intron | Splice region variant | LOW |
| splice_donor_region_variant | A sequence variant that falls in the region between the 3rd and 6th base after splice junction (5' end of intron) | Splice donor region variant | LOW |
| splice_polypyrimidine_tract_variant | A sequence variant that falls in the polypyrimidine tract at 3' end of intron between 17 and 3 bases from the end (acceptor -3 to acceptor -17) | Splice polypyrimidine tract variant | LOW |
| incomplete_terminal_codon_variant | A sequence variant where at least one base of the final codon of an incompletely annotated transcript is changed | Incomplete terminal codon variant | LOW |
| start_retained_variant | A sequence variant where at least one base in the start codon is changed, but the start remains | Start retained variant | LOW |
| stop_retained_variant | A sequence variant where at least one base in the terminator codon is changed, but the terminator remains | Stop retained variant | LOW |
| synonymous_variant | A sequence variant where there is no resulting change to the encoded amino acid | Synonymous variant | LOW |
| coding_sequence_variant | A sequence variant that changes the coding sequence | Coding sequence variant | MODIFIER |
| mature_miRNA_variant | A transcript variant located with the sequence of the mature miRNA | Mature miRNA variant | MODIFIER |
| 5_prime_UTR_variant | A UTR variant of the 5' UTR | 5 prime UTR variant | MODIFIER |
| 3_prime_UTR_variant | A UTR variant of the 3' UTR | 3 prime UTR variant | MODIFIER |
| non_coding_transcript_exon_variant | A sequence variant that changes non-coding exon sequence in a non-coding transcript | Non coding transcript exon variant | MODIFIER |
| intron_variant | A transcript variant occurring within an intron | Intron variant | MODIFIER |

*Table 1. Variant Effect Predictor (Ensembl VEP v111) variant effects classified by impact into 4 possible groups HIGH, MODERATE, LOW, MODIFIER. SO term is Sequence Ontology term. Note MODIFIER variants were pruned from final analysis.*

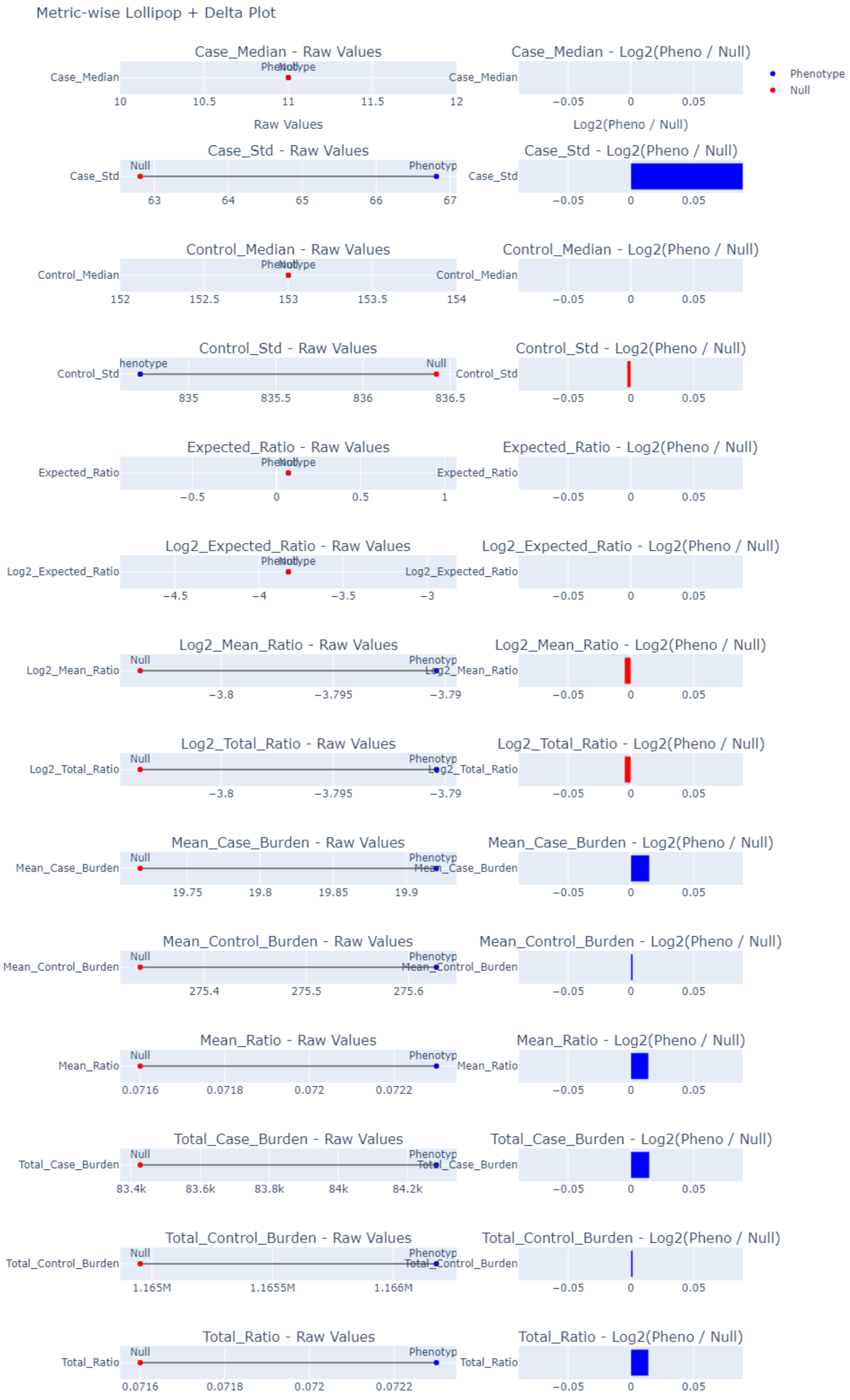


*Figure 1. Lollipop plot of cases to controls in pheno and null cohorts.*

| Metric | Phenotype | Null |
|---|---|---|
| Expected Case/Control Ratio | 0.071 | 0.071 |
| Log2(Expected Ratio) | -3.825 | -3.825 |
| Observed Total Burden Ratio | 0.072 | 0.072 |
| Log2(Total Ratio) | -3.79 | -3.804 |
| Mean Burden Ratio | 0.072 | 0.072 |
| Log2(Mean Ratio) | -3.79 | -3.804 |
| Total Case Burden | 84284 | 83426 |
| Total Control Burden | 1166179 | 1164952 |
| Per-sample Case Burden (mean) | 19.92 | 19.72 |
| Per-sample Case Burden (median) | 11 | 11 |
| Per-sample Case Burden (std) | 66.81 | 62.81 |
| Per-sample Control Burden (mean) | 275.63 | 275.34 |
| Per-sample Control Burden (median) | 153 | 153 |
| Per-sample Control Burden (std) | 834.72 | 836.42 |

*Table 2. Summary metrics of phenotype to null cohorts; cases (N=639) to controls (N=9059).*

| Variant Impact | MAF Range | Odds Ratio (OR) | P-value | 95% CI (log2 OR) |
|---|---|---|---|---|
| High | < 0.01% | 1.131 | <1×10-300 | 0.171 – 0.184 |
| | 0.01 – 0.1% | 0.96 | 4.62×10-85 | -0.064 – -0.053 |
| | 0.1 – 1.0% | 0.945 | 3.76×10-161 | -0.087 – -0.075 |
| Moderate | < 0.01% | 1.009 | 1.49×10-10 | 0.009 – 0.017 |
| | 0.01 – 0.1% | 0.989 | 2.73×10-15 | -0.020 – -0.012 |
| | 0.1 – 1.0% | 1.021 | 9.85×10-51 | 0.027 – 0.035 |
| Low | < 0.01% | 1.037 | 1.71×10-148 | 0.049 – 0.057 |
| | 0.01 – 0.1% | 1.015 | 4.43×10-27 | 0.018 – 0.026 |
| | 0.1 – 1.0% | 0.926 | <1×10-300 | -0.114 – -0.106 |
| Total Burden | All | 1.005 | 3.62×10-4 | 0.003 – 0.011 |

*Table 3. Global comparison of variant burden distributions using Firth's Logistic Regression.*

*The analysis compares the NMD phenotype cohort against the matched Null cohort across nine stratified bins and total gene burden. Odds Ratios (OR) > 1 indicate an enrichment of variants in the phenotype cohort. The strongest enrichment was observed in ultra-rare (<0.01%) High Impact variants ($OR=1.131, p<10^-300, confirming a baseline monogenic burden. Moderate Impact ultra-rare variants also showed significant enrichment (OR=1.009, p=1.49 * 10^-10, supporting their potential role in disease aetiology. Confidence Intervals (CI) represent the 95% interval for the log2(Odds Ratio).*

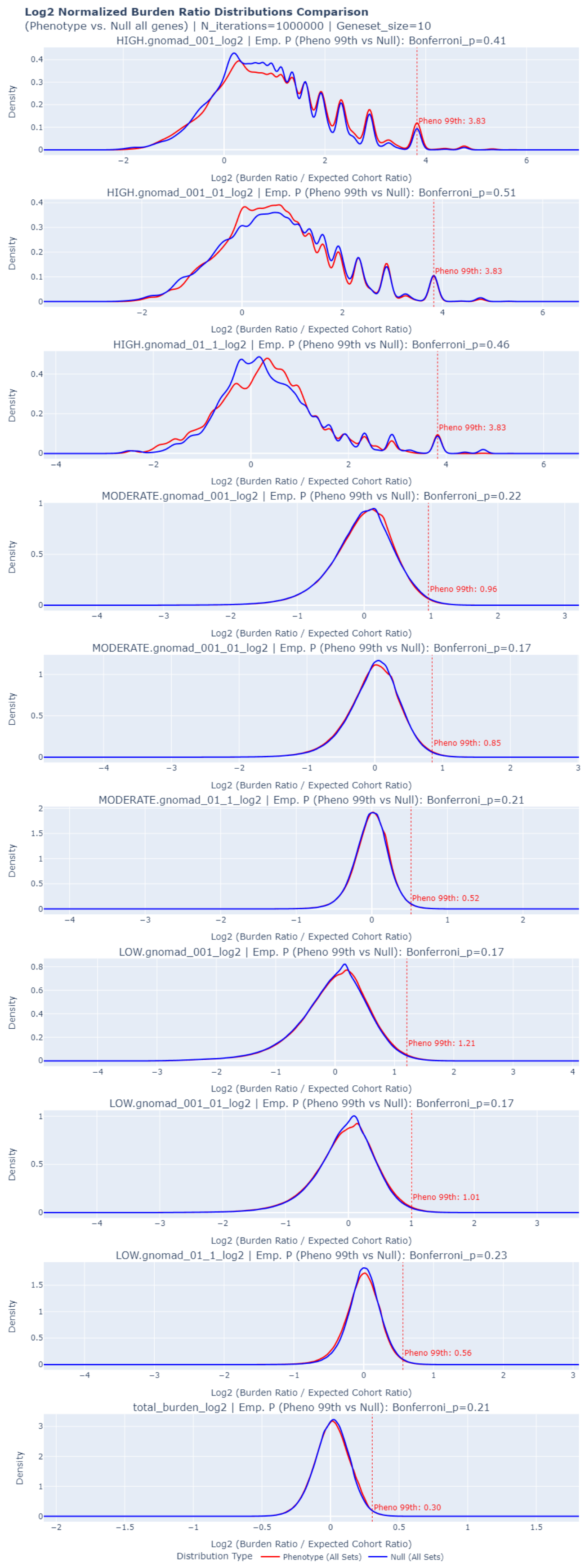


*Figure 2. Untruncated version of distributions of case/control ratios in 10 bins.*

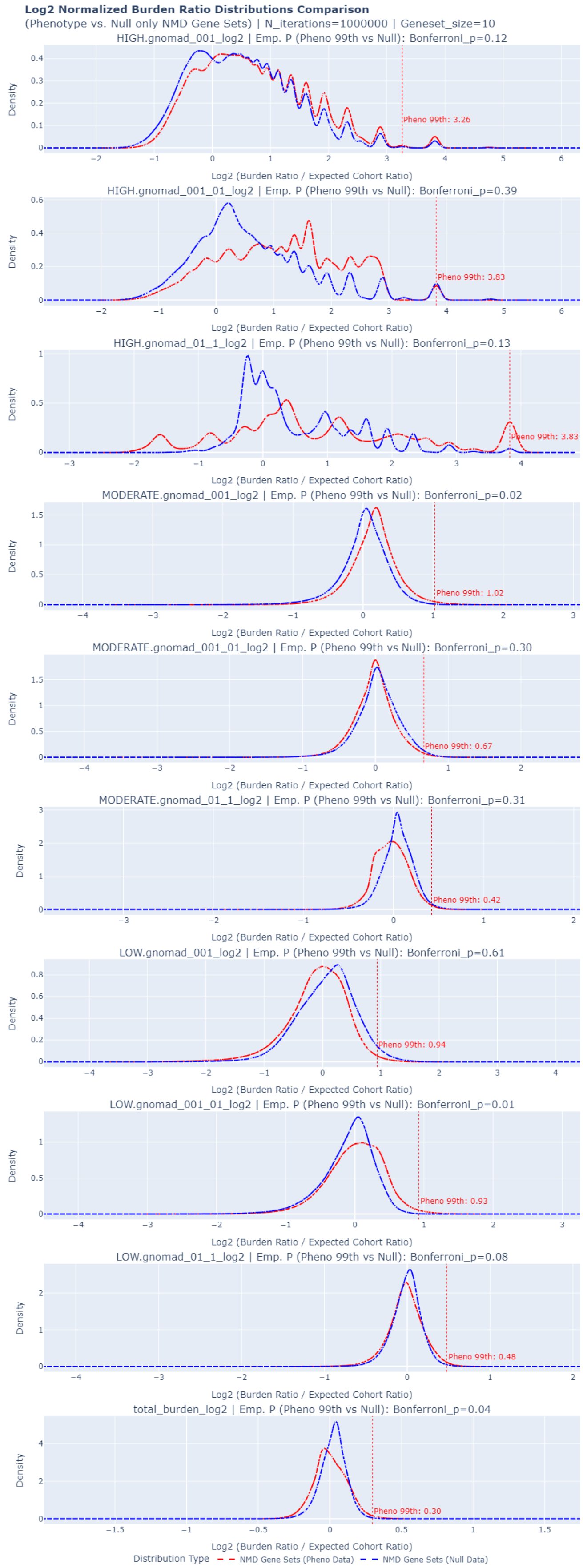


*Figure 3. Untruncated version of distributions of case/control ratios in 10 bins.*
